# Goldilocks and RNA: Where Mg^2+^ concentration is just right

**DOI:** 10.1101/2021.07.01.450745

**Authors:** Rebecca Guth-Metzler, Ahmad Mohyeldin Mohamed, Elizabeth T. Cowan, Ashleigh Henning, Chieri Ito, Moran Frenkel-Pinter, Roger M. Wartell, Jennifer B. Glass, Loren Dean Williams

## Abstract

Magnesium, the most abundant divalent cation in cells, catalyzes RNA cleavage but also promotes RNA folding. Because folding can protect RNA from cleavage, we predicted a “Goldilocks peak”, which is a local maximum in RNA lifetime at the Mg^2+^ concentration required for folding. Here we use simulation and experiment to discover an innate yet sophisticated mechanism of control of RNA lifetime. By simulation we characterized the RNA Goldilocks peak and its dependence on cleavage parameters and extent of folding. Supporting experiments with yeast tRNA^Phe^ and Tetrahymena ribozyme P4-P6 domains show that structured RNA can inhabit a Goldilocks peak *in vitro*. The Goldilocks peaks are tunable by differences in cleavage rate constants, Mg^2+^ binding cooperativity, and Mg^2+^ affinity. Broad ranges of those folding and cleavage parameters produce Goldilocks peaks of different intensities. Goldilocks behavior allows ultrafine control of RNA chemical lifetime, whereas non-folding RNAs do not display a Goldilocks peak. In sum, the effects of Mg^2+^ on RNA persistence are expected to be pleomorphic, both protecting and degrading RNA. In evolutionary context, Goldilocks behavior may have shaped RNA in an early Earth environment containing Mg^2+^ and other metals.

## Introduction

Universal biopolymers (DNA, RNA, protein, and carbohydrate) are ephemeral (1,2). Biopolymers hydrolyze spontaneously in aqueous media, degrading to monomers. In dilute aqueous solution, hydrolysis of biopolymers is always thermodynamically favored (3-5). However, very low rates of hydrolysis, due to kinetic trapping, allow biopolymers to persist for extended periods of time (1). Although RNA is especially labile (6), rates of RNA hydrolysis are modulated by many factors such as cations, RNA sequence, folding, temperature, and proteins (7-9).

Here we document and characterize Goldilocks behavior of RNAs, in which local maxima of chemical lifetimes are bounded by conditions of lower lifetimes **(Figure 1A)**. A Goldilocks landscape is a continuum of conditions, some of which are just right for RNA persistence. We predicted Goldilocks landscapes for RNA because Mg^2+^ directly increases RNA cleavage rates by one mechanism and indirectly decreases cleavage rates by a different mechanism. Mg^2+^, the most abundant divalent cation in cells (10), degrades RNA by catalyzing in-line attack of the 2’-oxygen on the backbone phosphate (7,11-16). Mg^2+^ can also protect against degradation, by facilitating RNA folding (17-22). The mechanism of protection involves converting conformationally flexible RNA to more structured RNA (23), which is less likely to adopt the geometry required for cleavage (24). A Goldilocks landscape arises when a given factor acts as a double-edged sword that differentially degrades and protects. Here, we investigated RNA Goldilocks behavior through simulation and experiment.

**Figure 1:**
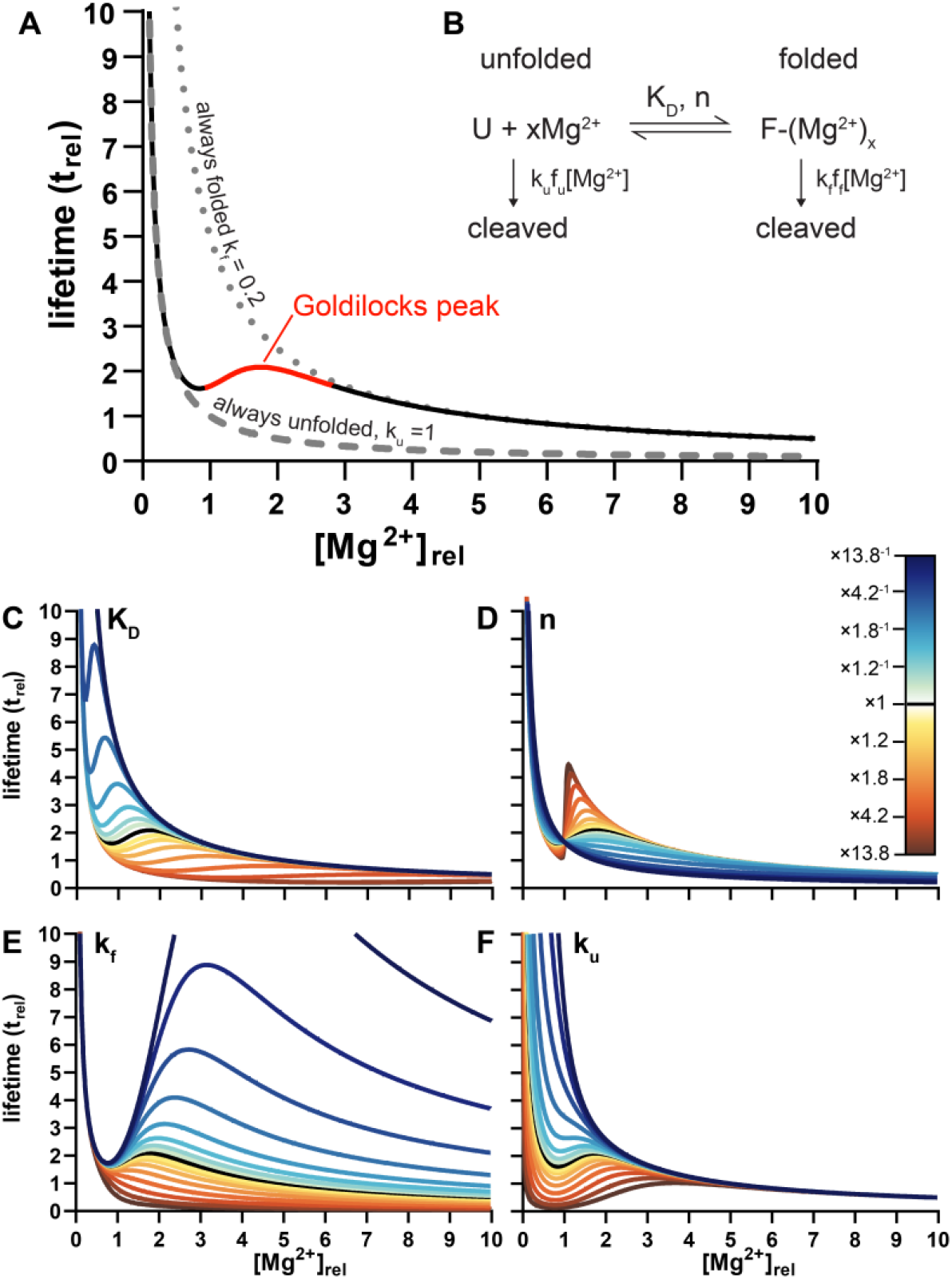
Goldilocks behavior of RNA is predicted by simulations. (A) Simulations reveal the influence of [Mg^2+^] on the chemical lifetime of an RNA that is cleaved more slowly in the folded state than in the unfolded state (black/red line). The Goldilocks peak is highlighted in red. The lifetime of a never folded RNA is shown by a dashed line (k_u_ = 1 t_rel_^-1^[Mg^2+^]_rel_^-1^). The lifetime of an always folded RNA is shown by a dotted line (k_f_ = 0.2 t_rel_^-1^[Mg^2+^]_rel_^-1^). An RNA that shifts between unfolded and folded states shifts between unfolded and folded lifetimes, to establish a Goldilocks peak. Goldilocks behavior requires conversion from unfolded to folded and a slower cleavage constant of folded vs. unfolded RNA (k_f_ < k_u_). (B) The two-state reaction mechanism. U is unfolded RNA and F is folded RNA. (C-F) Effects while other parameters are held constant of (C) K_D_ (D) n (E) k_f_, and (F) k_u_. Each parameter was varied by multiplication or division by 1+(0.1×2^i^) (i = 1, 2, 3, …8). For this representation, [Mg^2+^] was converted to [Mg^2+^]_rel_ where 1 [Mg^2+^]_rel_ = K_D_ = 0.022 mM Mg^2+^. Lifetime (t) was converted to t_rel_ where t_rel_ = 1 when [Mg^2+^]_rel_ =1 and the cleavage constant(s) are always 1 t_rel_^-1^[Mg^2+^]_rel_^-1^.

In simulations, Goldilocks behavior is observed under a broad variety of parameters that influence RNA folding and cleavage. Goldilocks landscapes are influenced by folding mechanism, Mg^2+^-dependency of folding, and folded and unfolded cleavage rate constants. Goldilocks landscapes are not accessible to RNAs that do not fold or unfold.

In experiments, Goldilocks behavior is observed for well-established model RNAs, yeast tRNA^Phe^ (25,26) and Tetrahymena ribozyme P4–P6 domain (27,28). An experimental comparison of tRNA and P4-P6 RNA indicates that the Goldilocks phenomena is retained even as the landscape is influenced by sequence and chemical modification. In experiments, Goldilocks peaks were observed where RNA is ∼95% folded; a control RNA that does not fold, rU_20_ (polyuridylic acid 20-mer), does not display Goldilocks behavior.

Goldilocks behavior of RNA suggests intrinsic sophistication, allowing ultrafine control of structure and chemical lifetime by a variety of inputs (1,29). RNA chemical lifetimes can be tuned by Mg^2+^-mediated shifts into and out of Goldilocks peaks and by remodeling Goldilocks landscapes via sequence and chemical modification. Goldilocks behavior of RNA is consistent with its selection in a primordial world of stringent and conflicting evolutionary demands.

## Materials and Methods

### Simulation of RNA lifetim

We mathematically modeled the effect of [Mg^2+^] on fraction of RNA folded and cleavage rate constants which together combine into an observed cleavage rate constant k_obs_. Lifetime is then the reciprocal of k_obs_.

For a two-state model, the fraction folded is f_f_ and the fraction unfolded is f_u_. We used the Hill equation to describe extent of folding (30), although any model that reasonably describes RNA folding can be used **(Eq. 1a and 1b)**:

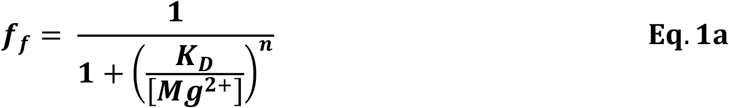

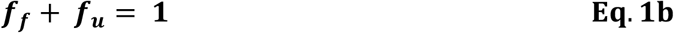

K_D_ is the [Mg^2+^] at the folding midpoint (21) and n is the Hill coefficient, which reflects the cooperativity of RNA folding.

The effects of Mg^2+^ on RNA cleavage are a second order phenomena in which the rate of cleavage depends on [RNA] and [Mg^2+^] (7,14). The observed pseudo-first order rate constant (k_obs_) is proportional to the second-order rate constant (k) and [Mg^2+^] **(Eq. 2)** (31):

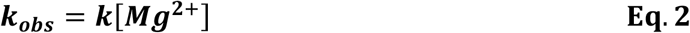

To model RNA lifetime with a two-state folding model, we used two cleavage rate constants: k_f_ for folded RNA and k_u_ for unfolded RNA. Folding offers protection from cleavage, and therefore k_f_ < k_u_. For an RNA that can occupy two states, k_obs_ is a convolution of cleavage contributions based on fractional occupancies and rate constants for each state **(Eq. 3**, which is an extension of **Eq. 2** with differential cleavage based on folding):

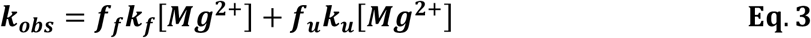

RNA lifetime is the reciprocal of the observed cleavage rate constant (k_obs_) **(Eq. 4)** (32):

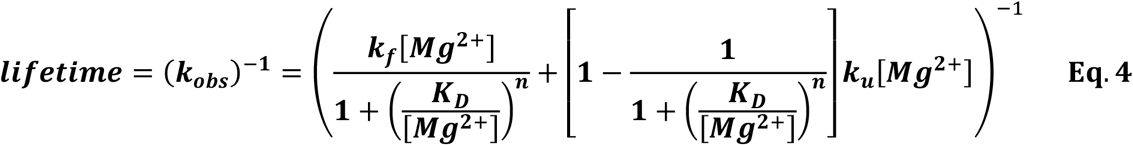

Selections of initial folding parameters to use in the lifetime equation, K_D_ = 0.022 mM Mg^2+^ and n = 4.1, were from previous work using native yeast tRNA^Phe^ (18). Initial values of relative rate constants were set to k_u_ = 1 and k_f_ = 0.2 t_rel_^-1^[Mg^2+^]_rel_^-1^, consistent with changes in RNA cleavage rates upon conversion of single strands to duplex (33,34). The software GraphPad Prism 8 was used to plot the simulations.

To model the contribution of folding intermediates to Goldilocks behavior, we modified the two-state model using the approach of Shelton et al. (18) **(Supp. Eq. 1)**. The transition from unfolded to intermediate is described by the terms K_D1_ and n_1_, and the transition from intermediate to fully folded is described by K_D2_ and n_2_. We initialized the three-state equation with K_D1_ at 1 [Mg^2+^]_rel_ and k_u_/k_f_ at 5. We set both n_1_ and n_2_ to 4.1 and K_D2_ for the second transition to 2 [Mg^2+^]_rel_. k_i_ was varied.

Our simulations and analysis of experimental data related to RNA Goldilocks phenomena assume that RNA folding is fast relative to cleavage. We assume both that Mg^2+^ binding to RNA is not rate-limiting and that cleavage operates over a fixed folding ensemble. These assumptions are based on experiment and theory. For folding, k = 10^4^ to 10^−4^ s^-1^ (35-42). For cleavage k = 10^−4^ to 10^−7^ s^-1^ (7,12,14,34). To either cleave or fold RNA, Mg^2+^ must first associate with the RNA. Diffuse association (or diffuse “binding”) of Mg^2+^ with RNA has a rate constant of k **≈** 10^10^ s^-1^, near the rate of diffusion itself (43), whereas specific binding, involving first shell coordination, has k **≈** 10^5^ s^-1^ (44,45).

### RNA Preparation

Yeast tRNA^Phe^ was purchased from Sigma-Aldrich (R4018). rU_20_ was a custom oligo purchased from Integrated DNA Technologies. T7-transcribed, stabilized P4-P6 RNA was produced as in Athavale et al. (46). Background Mg^2+^ was removed from the RNAs by dialysis in 180 mM NaCl and 50 mM HEPES buffer pH 7.1 using a 10 kDa MWCO filter.

### Circular Dichroism

Extent of folding was quantified by CD spectroscopy. A solution of 10 µM tRNA, 8 µM P4-P6 RNA, or 10 µM rU_20_ in 180 mM NaCl and 50 mM HEPES buffer pH 7.1 was added to a cuvette with a 0.1 cm path length. Spectra were accumulated on a Jasco J-815 spectropolarimeter with scan rate of 200 nm/min, bandwidth of 3 nm, and data pitch of 0.2 nm from 220 to 350 nm at 65°C. The RNAs were titrated with small volumes of concentrated known MgCl_2_ solutions. CD spectra were blank-subtracted and smoothed with a moving average. For yeast tRNA^Phe^ and P4-P6 RNA, fraction RNA folded was plotted using the theta value at the wavelength that maximized the difference between spectra (260 nm for tRNA and 260.6 nm for P4-P6). Theta values were corrected for the dilution that occurred upon Mg^2+^ addition then normalized.

### In-line cleavage of RNA

We approximated native ionic strength at 180 mM NaCl, which required elevated temperature (65°C) to observe Mg^2+^-dependent folding (47). Solutions of 10 µM yeast tRNA^Phe^, 3.8 µM P4-P6, or 10 µM rU_20_ in 180 mM NaCl, 50 mM HEPES buffer pH 7.1, and variable MgCl_2_ were incubated at 65°C for 48 hours. Reaction mixtures were separated by electrophoresis on 7% or 6% urea PAGE gels run at 120V for 1 hour and stained with SYBR Green II. Stained gels were digitized with an Azure 6000 or Typhoon FLA 9500 Imaging System. Band intensities were quantified using AzureSpot Analysis software to determine the amount of total intact RNA present.

### Sequencing and fragment analysis of P4-P6 RNA cleavage products

RNA species were analyzed by capillary electrophoresis (SeqStudio, Applied Biosystems) by following the manufacturer’s protocol and data was analyzed and aligned in MATLAB. In detail, a final concentration of 10 ng/µL uncleaved “fresh” P4-P6 RNA with 0.4 µM of FAM-labeled reverse transcription primer that binds to the 3’ end (5’-AGCTTGAACTGCATCCATATCAACA-3’, Integrated DNA Technologies), 5 mM DTT, 1X first strand buffer (Invitrogen), 0.5 mM each dNTP, and 2 mM of either ddATP, ddCTP, ddGTP, or ddTTP was reverse transcribed by SuperScript III (Invitrogen) to create fragments for sequencing. The P4-P6 RNA cleavage reactions containing 5 mM, 10 mM, and 15 mM Mg^2+^ were reverse transcribed in parallel with omission of the ddNTPs for identification of cleavage sites. 1 µL of each reverse transcription was combined with 1 µL GenefloTM 625 size standard ROX ladder (CHIMERx) and 20 µL HiDi (Applied Biosystems). Samples were resolved on a SequStudio instrument using the FragAnalysis run module.

## Results

### Simulations reveal Goldilocks behavior

We investigated RNA Goldilocks behavior for RNAs that fold in response to Mg^2+^. The simplest model **(Figure 1**) allows two states (folded and unfolded) and two cleavage rate constants; k_u_ is the cleavage rate constant of an unfolded RNA and k_f_ is the cleavage rate constant of a folded RNA. The observed rate constant shifts from k_u_ when the RNA is fully unfolded, to a weighted average of k_u_ and k_f_ when the RNA is partially folded, to k_f_ when the RNA is fully folded. This model allows a Goldilocks peak of chemical lifetime if k_u_ > k_f_. From the top of Goldilocks peak, RNA lifetime decreases if [Mg^2+^] is either increased or decreased. The folding transition in this model is governed by [Mg^2+^], K_D_ (the [Mg^2+^] at the folding midpoint and n (the Hill coefficient).

This simple two-state model predicts a Goldilocks landscape of chemical lifetime over a broad range of folding and cleavage parameters **(Figure 1C-F)**. K_D_ modulates the position of the Goldilocks peak on the [Mg^2+^] axis; RNAs that fold at lower [Mg^2+^] show a Goldilocks peak at lower [Mg^2+^]. The Hill coefficient n modulates the sharpness of the Goldilocks peak without a substantial change in its position; a larger n gives a sharper peak. k_u_ modifies the slope of the Goldilocks peak on the low [Mg^2+^] side, and k_f_ modifies the slope on the high [Mg^2+^] side. The ratio of k_u_ to k_f_ modulates the intensity of the peak. Goldilocks peaks are absent for RNAs that (i) do not fold, (ii) are always folded, (iii) do not change cleavage rate constant upon folding, or (iv) transition very gradually between differential cleavage realms with varying [Mg^2+^] **(Supp. Figure 1)**.

Here we define Goldilocks peak intensity as the ratio of the lifetime at the local maximum to the lifetime at the adjacent minimum. In simulation, positions of maxima and minima were determined by determining the simulated lifetime derivative across [Mg^2+^] and solving for [Mg^2+^] where slopes are zero. Returning each of those [Mg^2+^] values back into the original lifetime equation solves for the maximum and minimum used for Goldilocks peak intensity. In experiment, the low and high lifetime datapoints were used directly for the ratio.

### Goldilocks behavior in complex models

More realistic RNA folding mechanisms involve intermediate states (18). In a model with intermediates, each intermediate I is associated with specific cleavage rate constant k_i_ **(Figure 2A)**. The simulations reveal that the number, intensities, and proximities of Goldilocks peaks depend on the relative magnitudes of the rate constants and on locations of the folding transitions in Mg^2+^-space. For a three-state model with two transitions that are fully resolved in [Mg^2+^]-space, RNA can display two distinct Goldilocks peaks **(Figure 2B)**. When the transitions overlap in Mg^2+^-space, decreasing k_i_ tends to increase the intensity of the Goldilocks peak at the [Mg^2+^] where the intermediate population is maximum. Increasing k_i_ depresses lifetime at low [Mg^2+^] (peak position 1, **Figure 2B**) and shifts the Goldilocks peak to higher [Mg^2+^] (peak position 2, **Figure 2B**). An RNA with a folding intermediate that is cleaved more slowly than the fully folded state (k_i_ < k_f_) or cleaved more rapidly than the unfolded state (k_i_ > k_u_) it can display especially intense Goldilocks peaks. If an intermediate state has intermediate protection, the net Goldilocks peak is less intense than in the absence of an intermediate.

**Figure 2:**
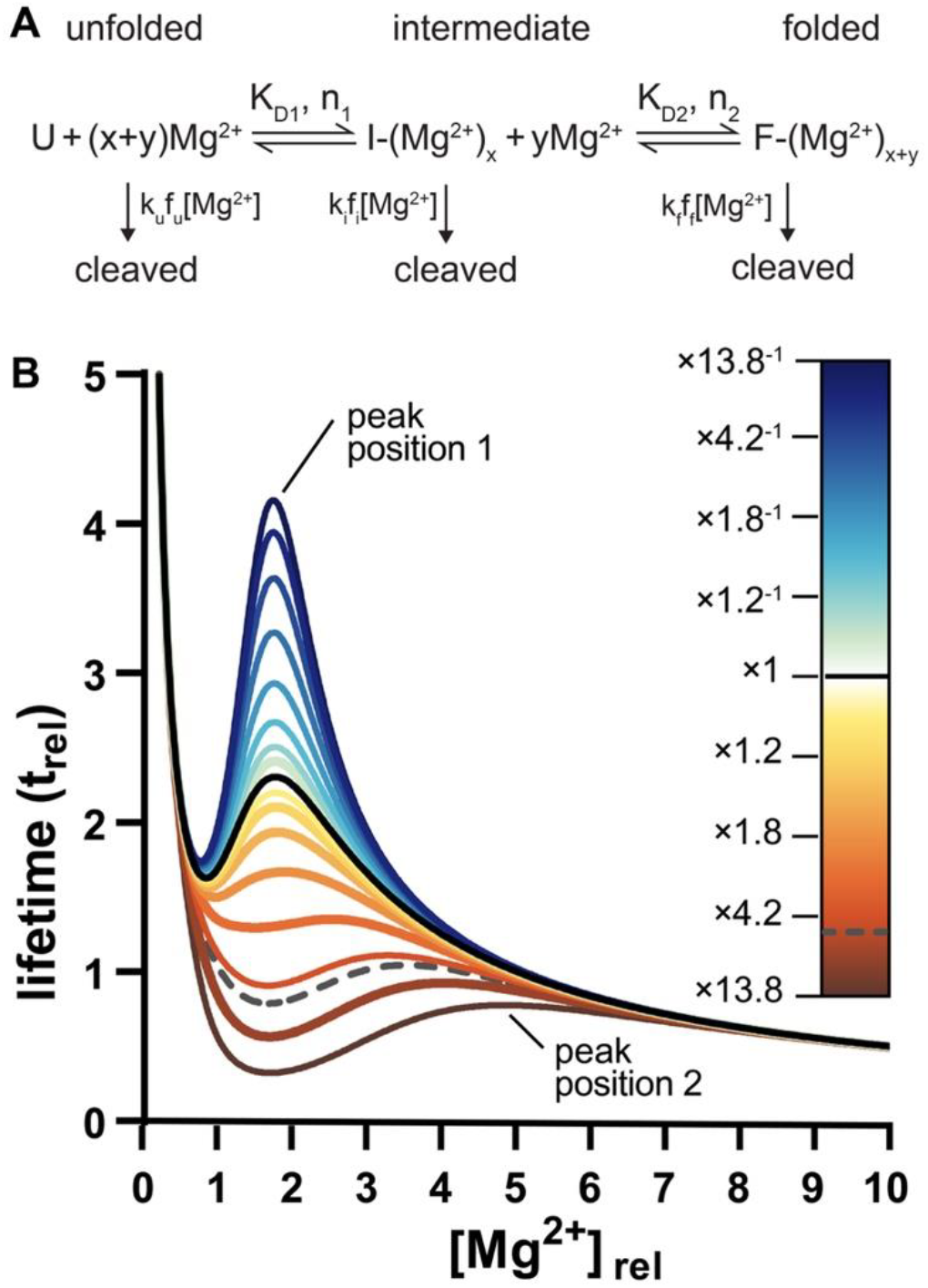
Complex folding models form Goldilocks peaks. (A) In a three-state mechanism, unfolded RNA converts by a first transition to an intermediate and by a second transition to fully folded. Unfolded RNA is cleaved with a rate constant k_u_, the intermediate is cleaved with a rate constant k_i_, and fully folded state is cleaved with a rate constant of k_f_. (B) In the simulation, k_i_ was varied relative to k_u_ while other parameters were fixed. The black line represents lifetimes when k_i_ = k_f_. The dashed line represents the lifetimes when k_i_ = k_u_. A k_i_ < k_f_ scenario favors an early Goldilocks peak and a k_i_ > k_u_ scenario favors a late Goldilocks peak.

### Goldilocks intensity

We show that RNA can inhabit a Goldilocks peak of RNA protection flanked by conditions of lability. The level of protection, given by Goldilocks peak intensity, depends on RNA properties. We defined Goldilocks peak intensity as the ratio of the peak maximum to minimum, i.e. the ratio of the protected lifetime to the labile lifetime. Using Goldilocks peak intensity, one can compare and rank various RNAs. We observed, in two-state simulations, that Goldilocks peak intensity increases with increased cooperativity of folding (n) or with increased extent of protection afforded by folding (decrease of k_f_ relative to k_u_) **(Figure 3A)**. We surveyed values of n and k_u_/k_f_ to create a Goldilocks intensity map **(Figure 3B)**. A k_u_/k_f_ = 3 with an n=4 produces a modest Goldilocks peak. Increasing either n or k_u_/k_f_ increases Goldilocks peak intensity. Lowering either disallows a Goldilocks peak unless there is a compensatory increase in the other.

**Figure 3:**
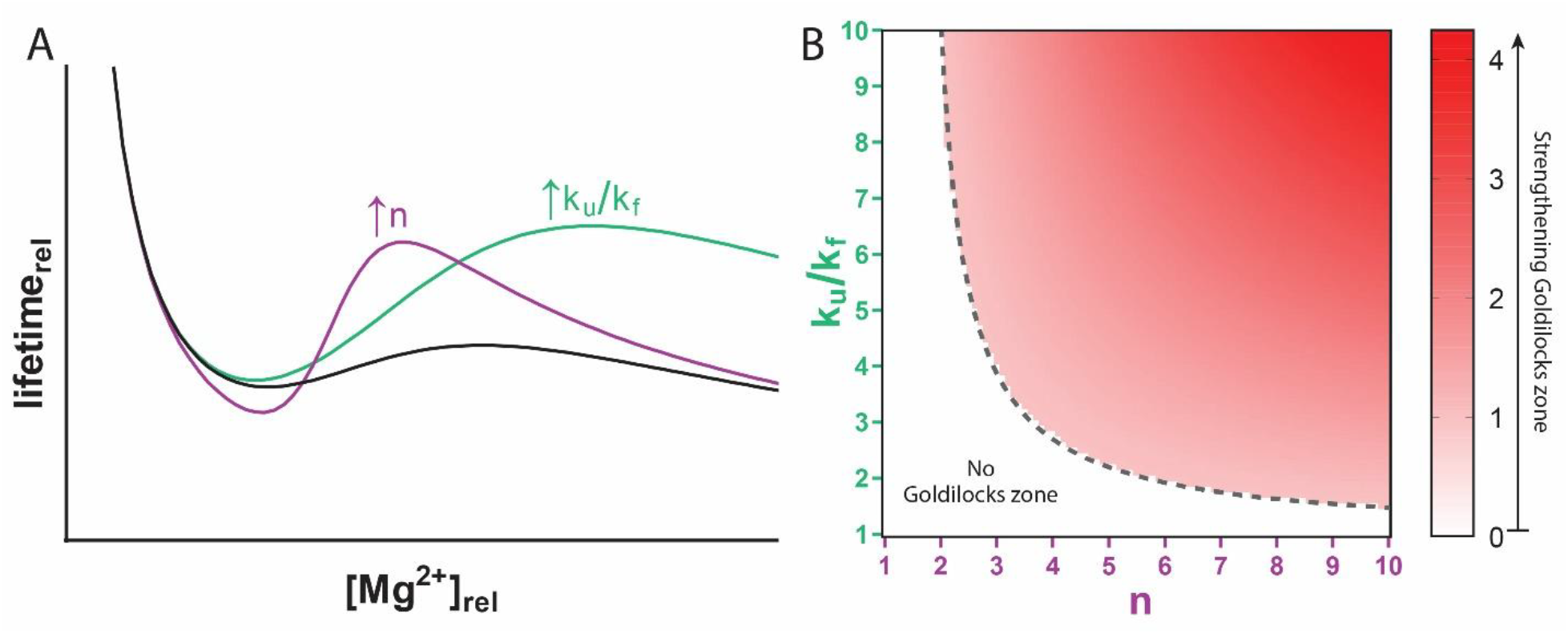
Goldilocks peak intensity increases with n and k_u_/k_f_ ratio. (A) A simulated Goldilocks peak (black) is enhanced by increasing the RNA’s cooperativity (n) or increasing the ratio of the unfolded to the folded cleavage rate constant (i.e. increasing protection of the folded state). (B) Surveying a range of rate constant ratios (k_u_/k_f_) across a range of n values shows that regions toward high k_u_/k_f_, n, or both have Goldilocks peaks and regions where both parameters are low do not.

### Experimental observation of a Goldilocks landscape of tRNA

To experimentally investigate Goldilocks behavior, we assayed both fraction folded and lifetime of yeast tRNA^Phe^ across a range of [Mg^2+^]. Circular Dichroism showed a clear cooperative folding transition with a [Mg^2+^] midpoint between 1 and 2 mM (**Figure 4 and Supplementary Data; Supp. Figure 2**). Random coil yeast tRNA^Phe^ folds to the native L-shaped structure upon addition of Mg^2+^ (35,48-53). Chemical lifetime of yeast tRNA^Phe^ showed a distinct Goldilocks peak near 3 mM Mg^2+^, where the tRNA was ∼95% folded **(Figure 4 and Supp. Figure 3)**. The tRNA lifetime was longer at 3 mM Mg^2+^ than at either 2.0 mM or at 3.5 mM Mg^2+^. In contrast, rU_20_, which does not fold, did not exhibit Goldilocks behavior **(Supp. Figure 4)**.

**Figure 4:**
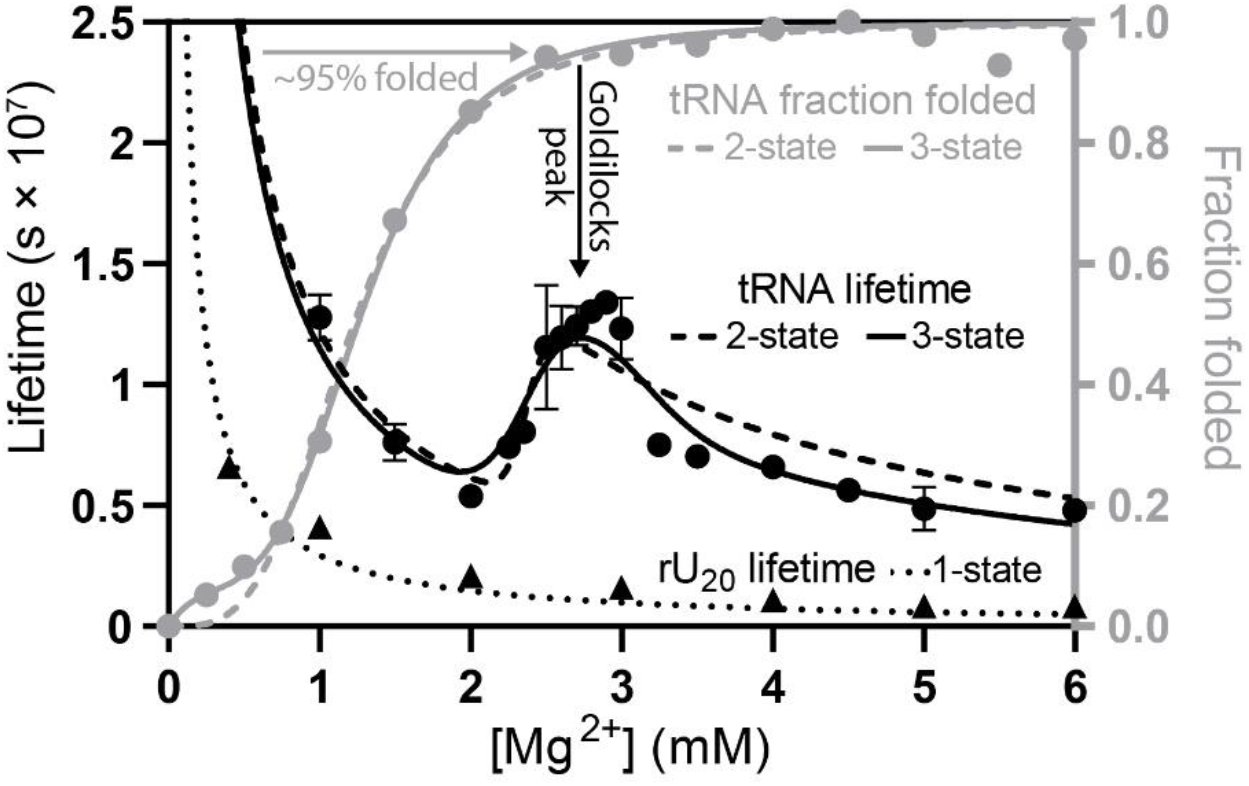
yeast tRNA_Phe_ shows a Goldilocks peak. Lifetimes of yeast tRNA^Phe^ (black circles) and rU_20_ (black triangles) were determined over a range of [Mg^2+^]. The yeast tRNA^Phe^ fraction folded (gray circles) was determined by CD. Experimental lifetimes were fit to two-state (black dashed) and three-state (black solid) folding models. Fraction folded was fit with two-state (gray dashed) and three-state (grey solid) Hill equation models. The three-state model better approximates the lifetime data, with a more intense Goldilocks peak than the two-state model. The tRNA Goldilocks peak is coincident with folding. rU_20_ lifetimes decrease monotonically with no Goldilocks peak (dotted black). rU_20_ did not show a folding transition **(Supp. Figure 4)**. Lifetimes were determined by quantification of intact RNA resolved by PAGE after 48 hours. All experiments were conducted in 180 mM NaCl, 50 mM HEPES pH 7.1 at 65°C, with variable [Mg^2+^]. Yeast tRNA^Phe^ lifetime error bars represent the standard deviation of five replicates. Folding and rU_20_ lifetime experiments used one replicate.

We compared the experimental lifetime data with predictions of our models (**Figure 4)**. The observed yeast tRNA^Phe^ Goldilocks landscape is reasonably fit by a two-state model. The fit and observed Goldilocks peaks are centered at the same [Mg^2+^]. The cleavage rate constant of the unfolded tRNA is predicted to be 2.7 times greater than that of the folded tRNA. However, the observed Goldilocks peak is sharper and more intense than predicted by the two-state model. A three-state model provides a better fit to the data, especially in the center of the Goldilocks peak. In the three-state model the cleavage rate constant for the unfolded RNA is predicted to be 3.2 times that of the intermediate and 2.2 times that of the fully folded tRNA (k_u_ > k_f_ > k_i_). The statistical significance of the improved fit of the three-state versus the two-state lifetime and folding models is indicated by residual errors **(Supp. Figure 5)**. Our results are consistent with previous observations of >2 states for folding of yeast tRNA^Phe^ (18,35,48,54). The fundamental conclusion here, which is the prediction and validation of Goldilocks behavior by RNA, is not dependent on the model.

Yeast tRNA^Phe^, with an intense Goldilocks peak, appears to fold via a protected intermediate. Prior work has shown that yeast tRNA^Phe^ is most compact in intermediate ionic strength (55-57) suggesting that the folding intermediate is more compact that the native state (however, see reference (58)).

### Experimental observation of a Goldilocks landscape of tetrahymena ribozyme P4-P6 domain RNA

P4-P6 RNA is a well-established model (27,28) that folds with increasing Mg^2+^ (59,60). The Mg^2+-^ dependence of P4-P6 RNA folding by CD **(Supp. Figure 6)** and chemical lifetime **(Supp. Figure 7 and Supplementary Data)** were determined **(Figure 5)** by the same methods and under the same conditions as for yeast tRNA^Phe^. P4-P6 RNA has a clear Goldilocks peak that is coincident with RNA folding.

**Figure 5:**
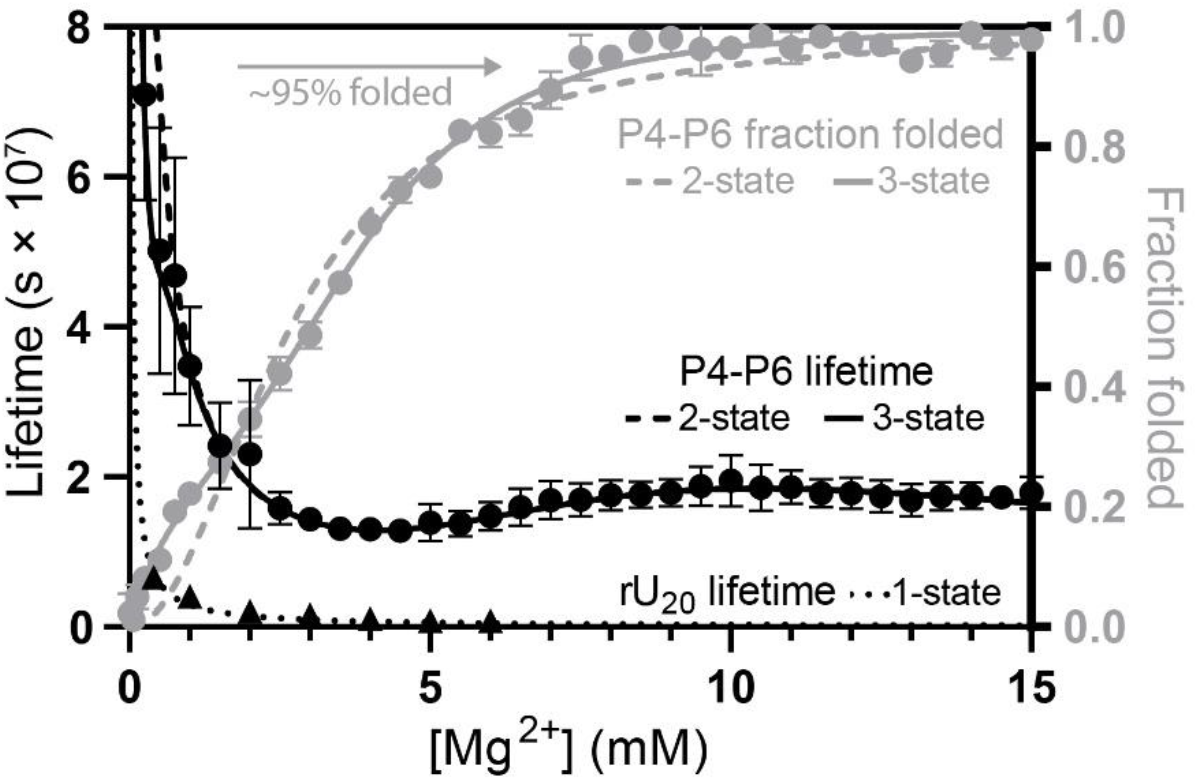
P4-P6 RNA shows a Goldilocks peak. P4-P6 RNA lifetime (black circles) shows a Goldilocks peak when the RNA is near-fully folded. The two-state (dashed line) and three-state (solid line) fits equally capture lifetime within the Goldilocks peak but the three-state fit better approximates lifetime at low [Mg^2+^]. Lifetime is normalized per phosphodiester bond. For a non-folding RNA comparison, rU_20_ lifetime (black triangles) is included and fit with a single state model (dotted line). P4-P6 RNA folding measured by CD (gray circles) is better approximated by a three-state fit (solid gray line) than a two-state fit (dashed gray line). Both folding and lifetime experiments were conducted in 180 mM NaCl, 50 mM HEPES pH 7.1 at 65°C with variable MgCl_2_. Error bars represent the standard deviations. P4-P6 RNA lifetime had four replicates, P4-P6 RNA folding had two replicates, and rU_20_ lifetime had one replicate.

The Goldilocks behavior of P4-P6 RNA is approximated by both the two-state or three-state models. Both models recreate the position in Mg^2+^-space and intensity of the single Goldilocks peak. The peak is produced by the intermediate to folded transition in the three-state model wherein k_i_ is 8 times k_f_. When constrained to two states, k_u_ is 22 times k_f_. A low [Mg^2+^] folding transition prior to the transition that forms the Goldilocks peak is approximated by the three-state model. The low [Mg^2+^] trend is captured only by the three-state model. The fit suggests that P4-P6 RNA folds by least three states, even though only two contribute to the Goldilocks peak. This conclusion is supported by residual plots for both lifetime and folding **(Supp. Figure 8)** and previous observations that P4-P6 RNA has more than two folding states (59,60).

### Comparison of yeast tRNA^Phe^ and P4-P6 RNA

Both RNAs display Goldilocks peaks in experiment and in simulation. Both RNAs are best fit to models with more than two states. For the tRNA the Goldilocks peak is sharp, with a half-height peak width of around 1 mM Mg^2+^. This level of cooperativity is a property of pronounced Goldilocks peaks. Conversely, P4-P6 RNA has low cooperativity, which is evident in broad Goldilocks peak, with a peak width at half-height of around 5 mM Mg^2+^. Goldilocks peak intensity for tRNA is greater (2.4) than for P4-P6 RNA (1.5). The fit parameters of each RNA are compared in **Table 1**.

**Table 1:**
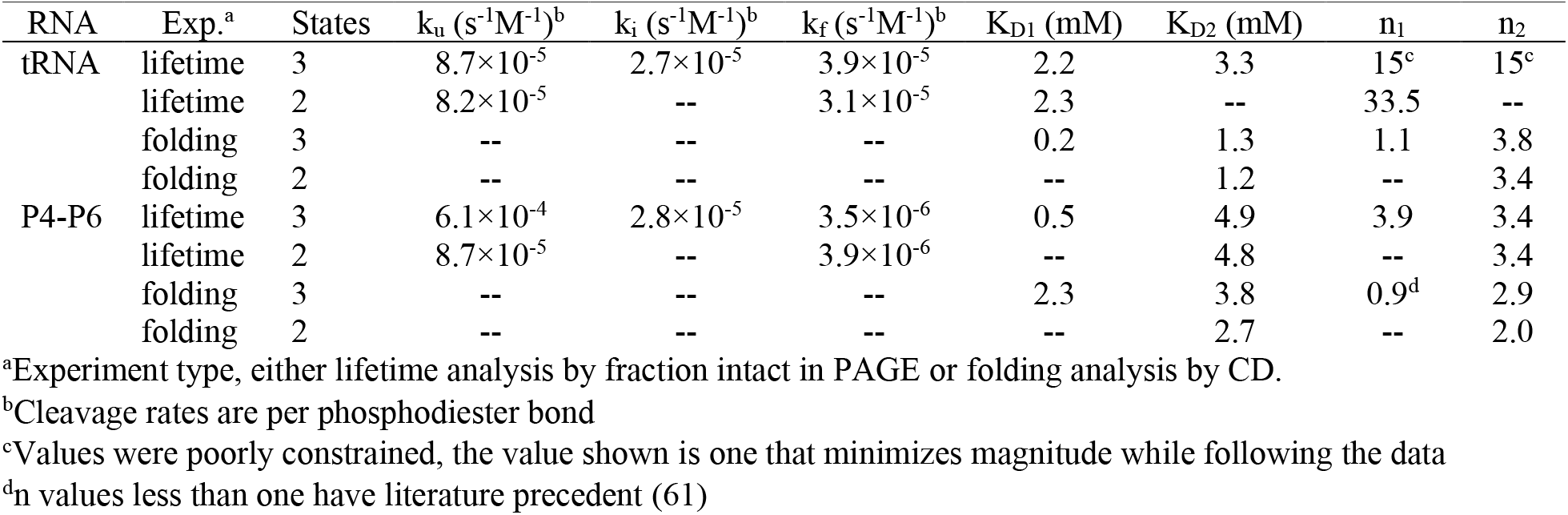
Fitted parameters for yeast tRNA_Phe_ and P4-P6 RNA.

| RNA | Exp.<sup>a</sup> | States | $k_u$ ($s^{-1}M^{-1}$)<sup>b</sup> | $k_i$ ($s^{-1}M^{-1}$)<sup>b</sup> | $k_f$ ($s^{-1}M^{-1}$)<sup>b</sup> | $K_{D1}$ (mM) | $K_{D2}$ (mM) | $n_1$ | $n_2$ |
| --- | --- | --- | --- | --- | --- | --- | --- | --- | --- |
| tRNA | lifetime | 3 | $8.7 \times 10^{-5}$ | $2.7 \times 10^{-5}$ | $3.9 \times 10^{-5}$ | 2.2 | 3.3 | 15<sup>c</sup> | 15<sup>c</sup> |
| | lifetime | 2 | $8.2 \times 10^{-5}$ | -- | $3.1 \times 10^{-5}$ | 2.3 | -- | 33.5 | -- |
|  | folding | 3 | -- | -- | -- | 0.2 | 1.3 | 1.1 | 3.8 |
|  | folding | 2 | -- | -- | -- | -- | 1.2 | -- | 3.4 |
| P4-P6 | lifetime | 3 | $6.1 \times 10^{-4}$ | $2.8 \times 10^{-5}$ | $3.5 \times 10^{-6}$ | 0.5 | 4.9 | 3.9 | 3.4 |
| | lifetime | 2 | $8.7 \times 10^{-5}$ | -- | $3.9 \times 10^{-6}$ | -- | 4.8 | -- | 3.4 |
|  | folding | 3 | -- | -- | -- | 2.3 | 3.8 | 0.9<sup>d</sup> | 2.9 |
|  | folding | 2 | -- | -- | -- | -- | 2.7 | -- | 2.0 |

<sup>a</sup>Experiment type, either lifetime analysis by fraction intact in PAGE or folding analysis by CD.
<sup>b</sup>Cleavage rates are per phosphodiester bond
<sup>c</sup>Values were poorly constrained, the value shown is one that minimizes magnitude while following the data
<sup>d</sup> $n$ values less than one have literature precedent (61)

### Site specificity of cleavage

To characterize Goldilocks phenomena at the level of single nucleotides, we quantified cleavage fragments of P4-P6 RNA using a sequencer. We examined the absolute extent and the site-specificity of cleavage under conditions of the Goldilocks peak (10 mM Mg^2+^), on the partially folded side of the peak (5 mM), and on the fully folded side of the peak (15 mM) **(Figure 6A-B)**. The extent of cleavage is less under the conditions of the Goldilocks peak (0.18) than in regions flanking the peak (0.21 pre-peak and 0.20 post-peak). The number of detectable cleavage sites and variety of cleavage intensities are greatest for the partially folded RNA (at 5 mM Mg^2+^). These sites overwhelm the few nucleotides that are highly cleaved in the folded state (strong negative Δcleavage). In the partially folded realm, double-stranded RNA shows more uniform extent of cleavage than unpaired RNA (62). Variability decreases when the RNA fully folds, where the conformations of essentially all nucleotides become fixed. Here, Δcleavage for 5 mM to 10 mM Mg^2+^ is most variable, especially in bulge and loop-forming regions. When increasing from 10 mM to 15 mM Mg^2+^ a few cleavage hot spots emerge in unpaired regions, where the RNA is most susceptible to cleavage (24).

**Figure 6:**
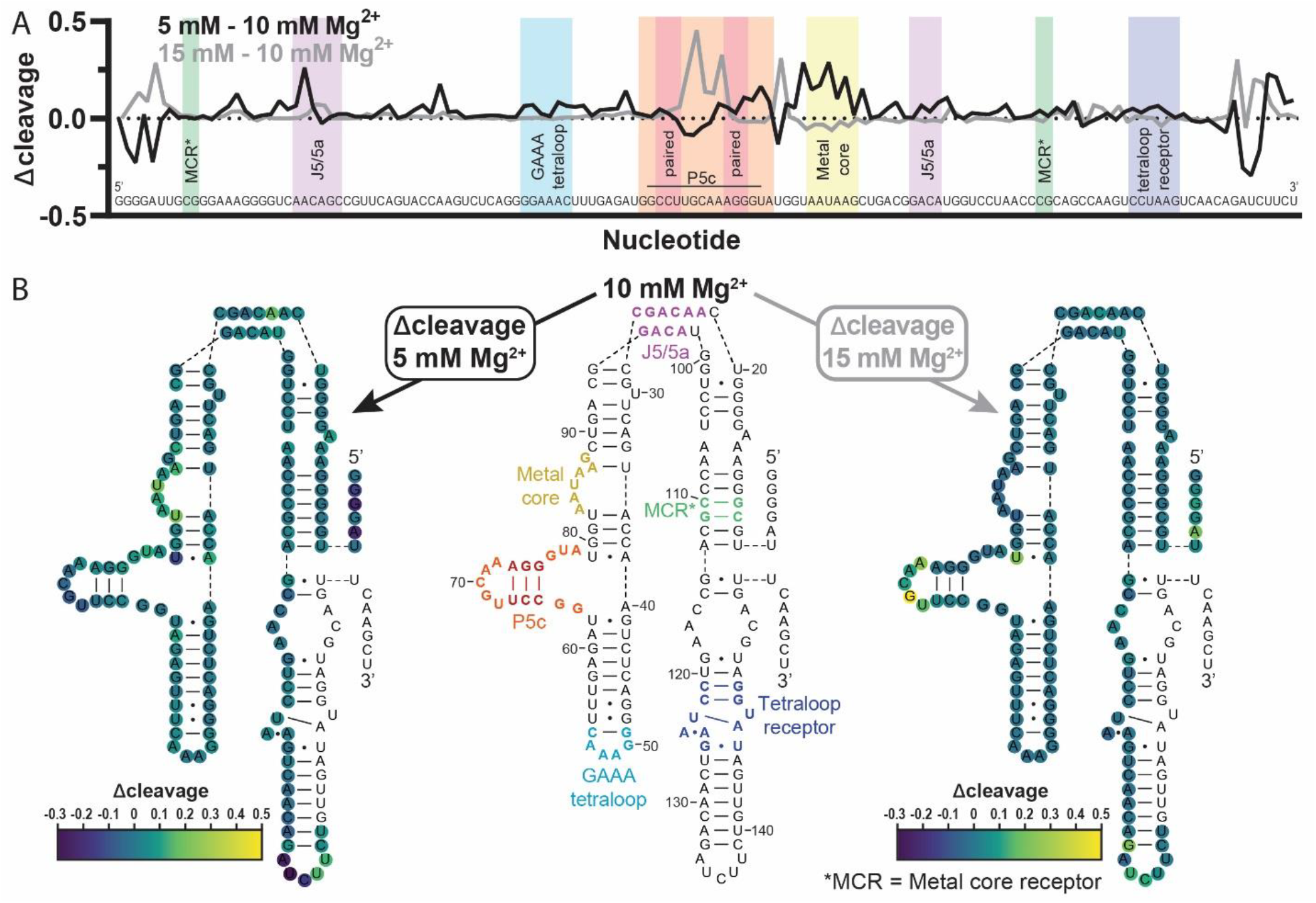
Differences in site specificity of cleavage with changing [Mg_2+_]. (A) Δcleavage reports differences in cleavage between P4-P6 RNA at the Goldilocks peak (10 mM Mg^2+^) and at 5 or 15 mM Mg^2+^. Δcleavage indicated highest lifetime at the Goldilocks peak. (B) Superimposition of cleavage data onto the secondary structure of P4-P6 RNA shows that hot spots for cleavage (yellows and light greens) or protection (dark blue) relative to P4-P6 RNA at 10 mM Mg^2+^ occur at loops and bulges in the RNA. Important structural features are provided as in Bisaria 2016 (64). Each Δcleavage value is the mean of two replicates.

In P4-P6 RNA, the Mg^2+^ core does not appear to be folded at 5 mM Mg^2+^, as indicated by extent of cleavage **(Figure 6A-B)**. The metal core folds and is protected at 10 mM Mg^2+^. This region is a representation of the double-edged sword of Mg^2+^; by associating with Mg^2+^ the RNA is protected from Mg^2+^. As expected (63), by its low reactivity relative to other loop or bulge regions, the GAAA tetraloop appears to be fully folded by 5 mM Mg^2+^.

The sequencing data indicate that even though the low resolution P4-P6 RNA PAGE banding patterns remain reasonably constant, relative extent of cleavage at various sites does in fact change. Although the yeast tRNA^Phe^ banding pattern appeared uniform across [Mg^2+^] in gels, we assume significant differences in locations of cleavage upon folding. Site-specific analysis of yeast tRNA^Phe^ was not possible with our method because of the RNA base modifications.

## Discussion

By simulation and experiment we validated a Goldilocks model of RNA. Local maxima in lifetime are flanked by conditions of greater lability. RNAs can resist Mg^2+^-mediated cleavage when Mg^2+^ folds the RNA. Increasing [Mg^2+^] beyond the folding threshold increases Mg^2+^-mediated cleavage. We use Goldilocks model framework to explain how lifetime landscapes are modulated by specific characteristics of diverse RNAs. We predict that Goldilocks landscapes are modulated by monovalent cation concentrations, type of divalent cation, RNA sequence and modification, protein and ligand association, and temperature. RNA that cannot fold or unfold cannot access a Goldilocks peak. Self-cleaving ribozymes are exempt from Goldilocks behavior because their folding increases rates of cleavage.

### Goldilocks landscapes

RNA is responsive to [Mg^2+^] on a landscape that is modulated by RNA sequence and chemical modification. The number, intensity, profile, and position in [Mg^2+^]-space of Goldilocks peaks depends on RNA sequence and chemical modification. The position of a Goldilocks peak in [Mg^2+^] space is determined primarily by the affinity of the folded RNA for Mg^2+^. A smaller K_D_ shifts the Goldilocks peak to lower [Mg^2+^].

The Goldilocks landscape of RNA should extend beyond Mg^2+^ to species such as Fe^2+^, which also promotes both RNA folding and cleavage (46,65). Even farther, the general principles of Goldilocks behavior can be applied to any agent that has differential “positive” and “negative” effect on biopolymer lifetime. For example, protein is cleaved by hydrolysis (66). Protein folding decreases rates of hydrolysis (1) and is promoted by high water activity (67). This model predicts water-defined Goldilocks phenomena for proteins.

### The Goldilocks peak in vivo

Our simulations anticipate some RNAs *in vivo* may inhabit Goldilocks peaks. Free Mg^2+^ *in vivo* is near 1 mM (68,69) **(Table S1**). An RNA hairpin ribozyme used as a model is mostly folded at 1 mM Mg^2+^ in molecularly crowded conditions mimicking the cell (70). If the minimal [Mg^2+^] required for folding in vivo coincides with the in vivo [Mg^2+^] RNA may occupy a Goldilocks peak. More specific in vivo conclusions remain unresolved thus far because of differences in in vitro and in vivo conditions and limitations in manipulating in vivo [Mg^2+^] (71). Goldilocks landscapes remain to be evaluated in the context of protein and ligand binding and RNase hydrolysis in the cell. However, Goldilocks behavior could explain in part why cells invest in careful maintenance of Mg^2+^ homeostasis (72). It seems likely that a narrow range of [Mg^2+^] prolongs specific RNA lifetimes *in vivo*.

### Goldilocks and Origins of Life

RNA can shift from dangerous spaces to safe spaces. This finely controlled metastability, including access to Goldilocks peaks of protection, are likely to be imprints of evolutionary processes leading to the emergence of RNA on the ancient Earth. In this scenario, acute control of lifetime was a selected trait (73); and backbone structure, base modifications, and sequences, all of which modulate Goldilocks landscapes, are the products of selection.

## Availability

All data are available through NAR online.

## Supplementary Data

Supplementary Data are available at NAR online.

## Acknowledgment

We thank Drs. Jonathan B. Chaires, Nicholas V. Hud, and Ramanarayanan Krishnamurthy for helpful discussions.

## Funding

This work was supported by the National Aeronautics and Space Administration Astrobiology Program Center for the Origins of life 80NSSC18K1139.

## Notes

### Competing Interest Statement

The authors have declared no competing interest.

### Summary of Updates

More simulations and experiments with an additional RNA were added and supported the existing model. The text was updated for these additions and restructured accordingly for the evolution of the project.

