## Supplementary Information for "Goldilocks and RNA: Where Mg^2+^ concentration is just right"

**This PDF includes:**

Supplemental Equation 1
Supplemental Table 1
Supplemental Figures 1-8
Supplemental References

**Supplemental Equation 1:** Lifetime of RNA with three states.

$$lifetime = \frac{1}{k_{obs}} = \frac{1}{[Mg^{2+}]} \times \frac{1 + \left(\frac{[Mg^{2+}]}{K_{D1}}\right)^{n1} + \left(\frac{[Mg^{2+}]}{K_{D1}}\right)^{n1}\left(\frac{[Mg^{2+}]}{K_{D2}}\right)^{n2}}{k_u + k_i\left(\frac{[Mg^{2+}]}{K_{D1}}\right)^{n1} + k_f\left(\frac{[Mg^{2+}]}{K_{D1}}\right)^{n1}\left(\frac{[Mg^{2+}]}{K_{D2}}\right)^{n2}}$$

The overall lifetime can also be written simply as the proportional contributions of the lifetimes of RNA in each state, where the fraction of RNA in each state adds to one.

$$lifetime = f_u lifetime_u + f_i lifetime_i + f_f lifetime_f$$

$$f_u + f_i + f_f = 1$$

**Supplemental Table 1:** Cytosolic free $Mg^{2+}$ (non-complexed and unbound) concentrations in bacteria and eukarya. References (1, 2) are compiled from previous studies.

| **Species and cell type** | **Free $Mg^{2+}$ (mM)** | **Reference** |
|---|---|---|
| *Escherichia coli* (bacterium) | 0.8±0.2 | (3) |
| *Salmonella enterica* (bacterium) | 0.9-1.5 | (4) |
| *Saccharomyces cerevisiae* (yeast) | 0.9-2.0 | (5) |
| *Penicillium chrysogenum* (fungi) | 0.4-0.8 | (5) |
| *Endomyces magnusii* (fungi) | 0.4-1.6 | (5) |
| *Xenopus laevis* (giant squid) | 0.3 | (6) |
| Skeletal muscle (human, mouse, frog, rat) | 0.6-1.3 | (1, 7) |
| Cardiac myocytes (chicken, rat, guinea pig) | 0.5-1.1 | (1, 2) |
| Smooth muscle cells (rabbit, rat, guinea pig) | 0.3-1.0 | (1) |
| Neuronal cells (human) | 0.7-1.0 | (1) |
| Exocrine cells (rat, lymphocytes) | 0.2-0.4 | (1) |
| Adrenal cells (rat) | 0.5-0.9 | (8) |
| Red blood cells (human) | 0.3-1.9 | (1, 7) |
| Hepatocytes | 0.3-0.7 | (1) |

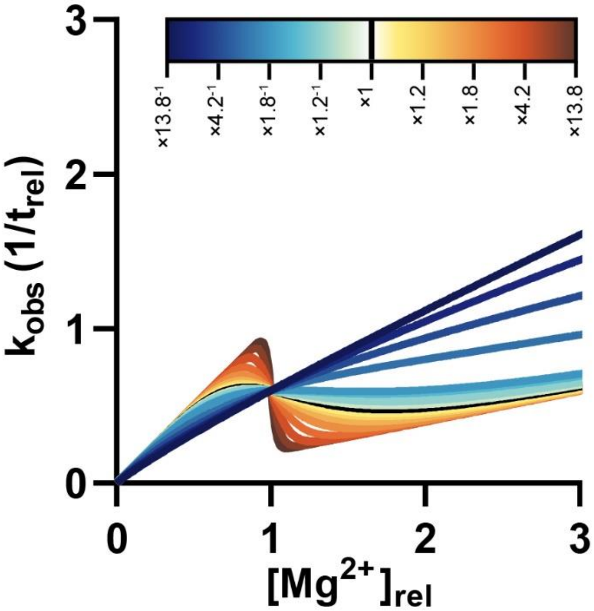


**Figure S1: The Goldilocks peak requires cooperative binding of $Mg^{2+}$ to RNA.** $k_{obs}$ is the reciprocal of lifetime. When lifetime rises $k_{obs}$ decreases and vice versa. The $k_{obs}$ vs. [$Mg^{2+}$] graph has the advantage of straight trends and easy visualization. For very low n there is not an inflection of $k_{obs}$ with increasing [$Mg^{2+}$] (darker blue lines). So, lifetime does not increase, and no Goldilocks peak occurs. The same parameters and scale were used as in **Figure 1D**, where the initial n is equal to 4.1 (black line) and the various colors have n values that are the initial n multiplied by the indicated factor.

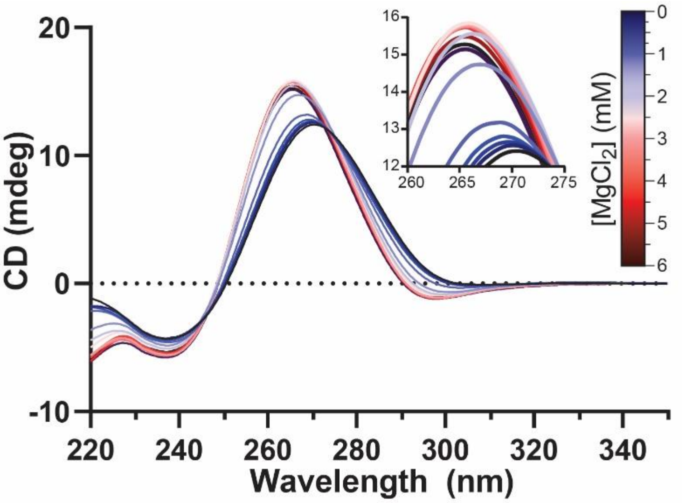


**Figure S2: tRNA folds with increasing [$Mg^{2+}$].** Yeast $tRNA^{Phe}$ shows a change in CD spectra upon addition of $MgCl_2$ indicating a $Mg^{2+}$-induced structural transition. To extract the fraction of full-length RNA from the spectra, the theta values at 260 nm were corrected for dilution upon $MgCl_2$ addition then plotted against the [$Mg^{2+}$]. The experiment was run in 180 mM NaCl, 50 mM HEPES pH 7.1 at 65°C with variable $MgCl_2$.

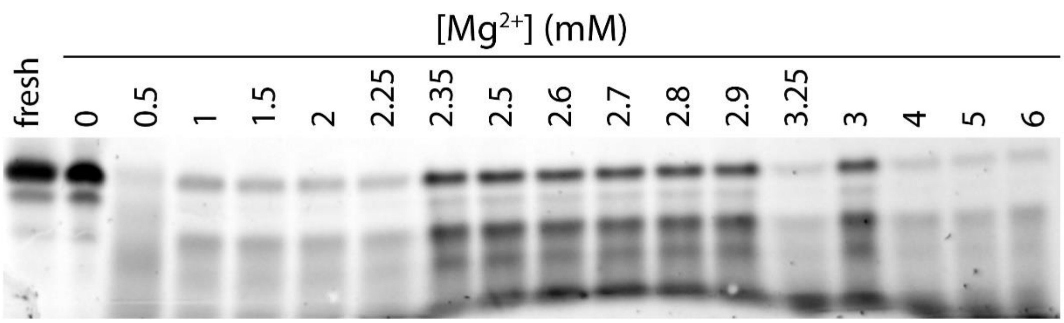


**Figure S3: the fraction of intact tRNA has a multiphasic response to increasing [$Mg^{2+}$].** A representative gel of yeast $tRNA^{Phe}$ after no reaction (fresh) or 48 hours of cleavage shows that the fraction of intact RNA first decreases, then increases, then decreases again when exposed to increasing [$Mg^{2+}$]. Quantification in our gels of the top band, i.e. the full-length band, produced our fraction intact and lifetime data. The re-emergence of the full-length band does not depend on position in the gel, shown by loading some samples out of order.

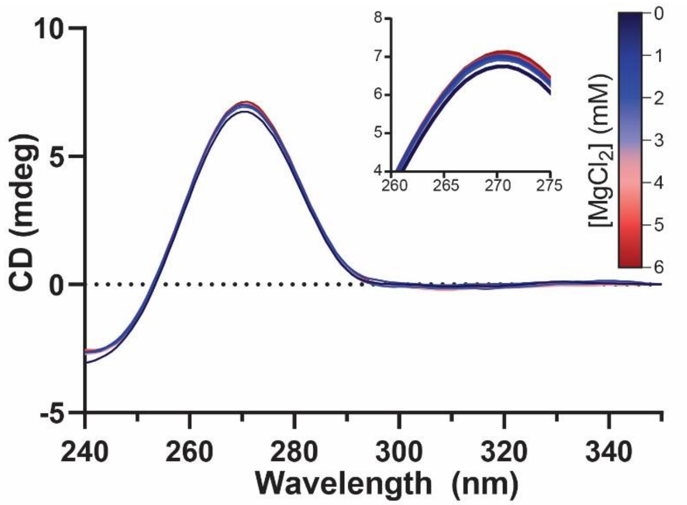


**Figure S4: $rU_{20}$ does not fold.** $rU_{20}$ does not show a change in CD spectra upon addition of $MgCl_2$ as expected for a non-folding RNA. The experiment was run in 180 mM NaCl, 50 mM HEPES pH 7.1 at 65°C with variable $MgCl_2$.

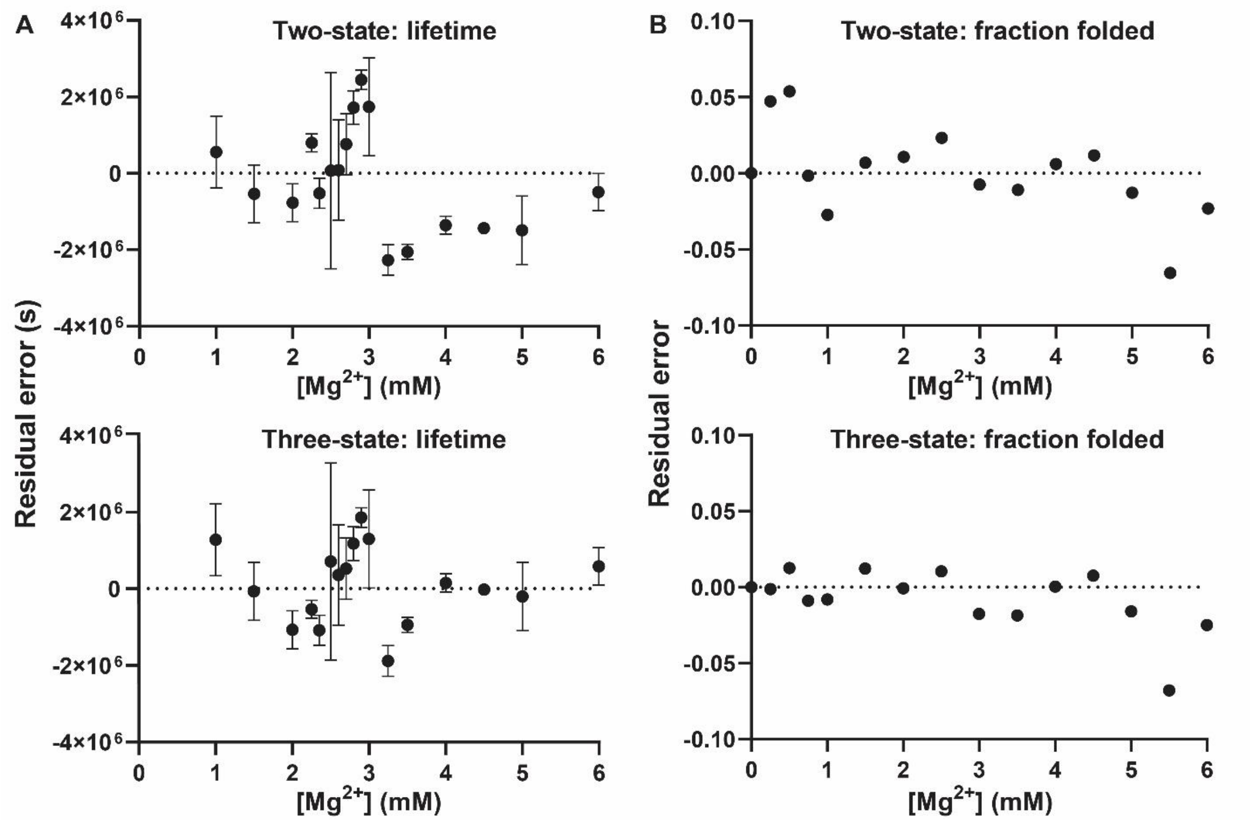


**Figure S5: The tRNA lifetime and folding residual errors are minimized by the three-state model.** (A) For the two-state model the lifetime errors are large and non-random; all the mean errors are positive between 2.8 and 3.0 mM $Mg^{2+}$ and are negative for all values of [$Mg^{2+}$] greater than 3.0. For the three-state model the errors appear to be more randomly distributed around zero. (B) Errors in folding are minimized by the three-state model below 2.5 mM $Mg^{2+}$, showing better approximation of an early transition state.

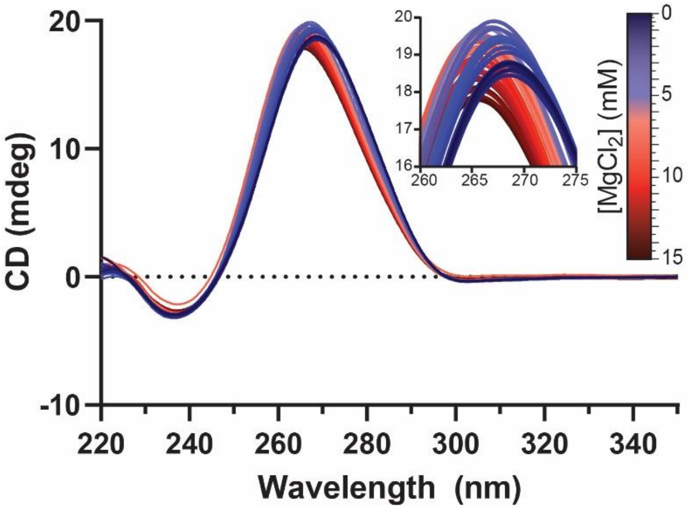


**Figure S6: P4-P6 RNA folds with increasing [$Mg^{2+}$].** P4-P6 RNA shows a change in CD spectra upon addition of $MgCl_2$ indicating a $Mg^{2+}$-induced structural transition. The initial rise in peak height is the $Mg^{2+}$ folding response, which is followed by a peak decrease, the effect of dilution when adding $Mg^{2+}$ that becomes apparent when the RNA has completed folding. To extract the fraction of full-length RNA from the spectra, the theta values at 260.6 nm were corrected for dilution then plotted against the [$Mg^{2+}$]. The experiment was run in 180 mM NaCl, 50 mM HEPES pH 7.1 at 65°C with variable $MgCl_2$.

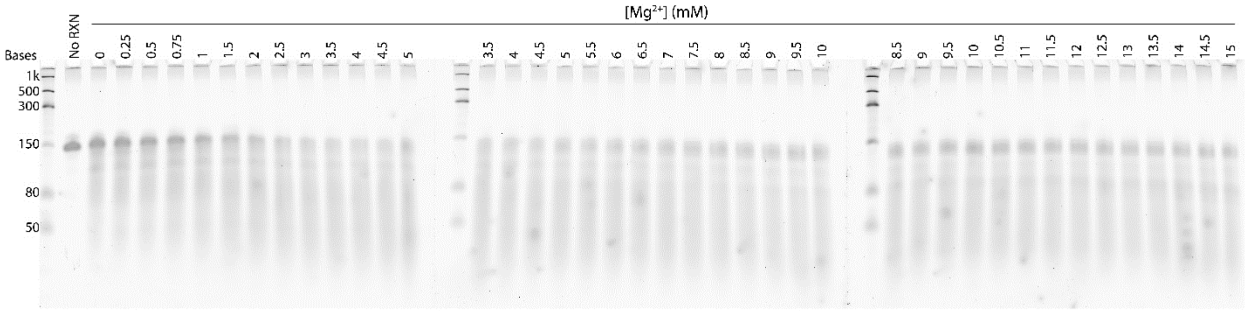


**Figure S7: the fraction of intact P4-P6 RNA has a multiphasic response to increasing [$Mg^{2+}$].** A representative gel of P4-P6 RNA after 48 hours of cleavage shows that the fraction of intact RNA first decreases, then increases, then decreases again when exposed to increasing [$Mg^{2+}$]. A surplus of prepared RNA allowed a single mixture to be added to up two gels. In this way each gel was made to share four samples with its neighboring gel allowing for normalization between gels and for the running of the large number of samples. Additionally, running and then imaging was performed simultaneously for the three gels.

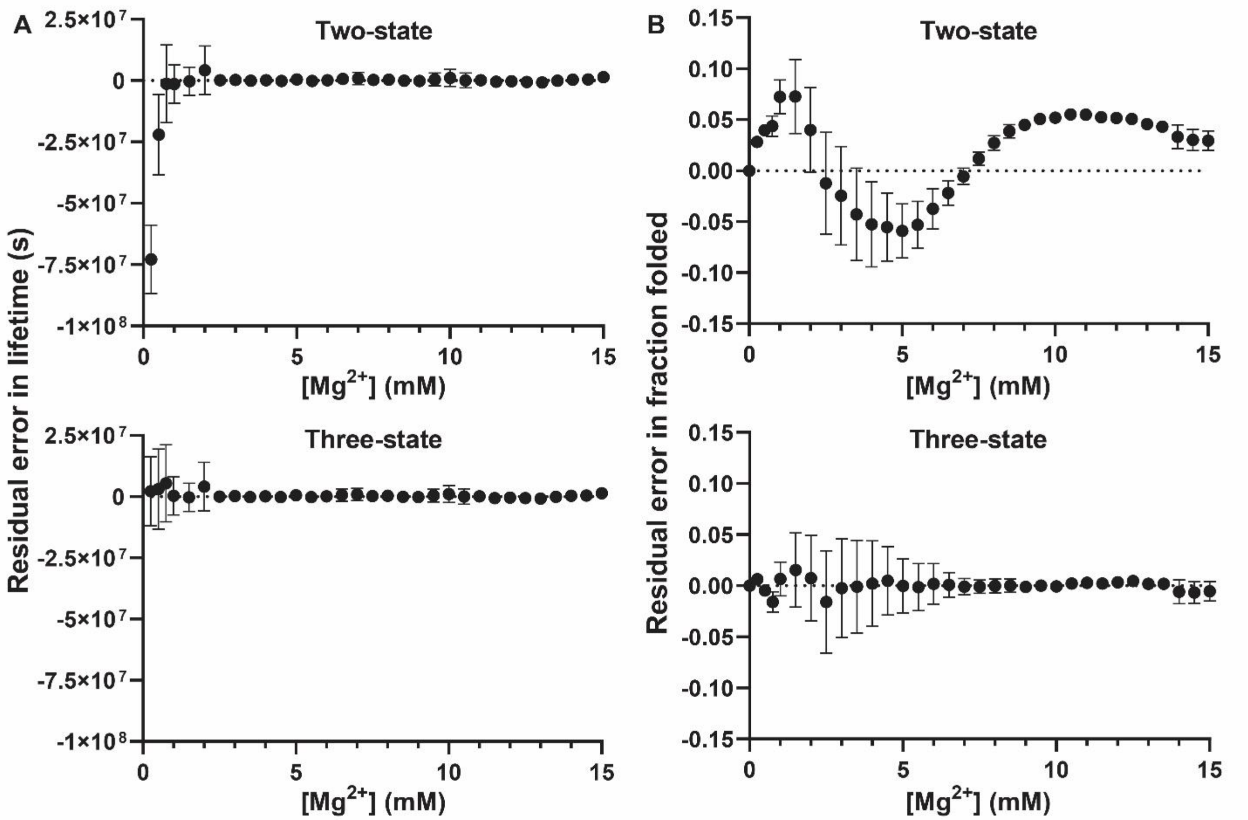


**Figure S8: P4-P6 RNA has an apparent transition to an intermediate folding state at low [$Mg^{2+}$].** (A) Lifetime residuals between two- and three-state models only differ at low [$Mg^{2+}$] where only the three-state model captures the data of an apparent early folding state. (B) Residual error in the two-state model systematically weaves above and below zero while the three-state difference is smaller and spreads randomly near zero. The three-state fit is a more accurate representation of the folding of P4-P6 RNA again showing an intermediate state between unfolded and folded.
